# PABA Release from Chitosan-PCL with Induced Electric Current

**DOI:** 10.1101/377481

**Authors:** Jennifer M. Miller, Roche C. de Guzman

**Affiliations:** Bioengineering Program, Department of Engineering, Hofstra University, Hempstead, New York, USA

**Keywords:** Drug Delivery, Chitosan, PCL, Biomaterial Scaffolds, PABA, Electrical Stimulation, Electrostatic Interaction

## Abstract

Controlled drug delivery systems such as the stimulation-based biomaterial scaffolds for sequestration and release of drugs offer safety and regulated therapeutic approach. In this study, the drug: *para*-aminobenzoic acid (PABA) was absorbed into a crosslinked chitosan and poly(caprolactone) (PCL) hydrogel and its release kinetics quantified under different conditions. It was experimentally-observed that the higher the pH (or the more basic the pH), the slower the PABA saturation release trended over time. At the acidic environment of pH 4, PABA was released the fastest, and enhanced by the degradation of chitosan-PCL gel. When a constant electric current of 0.6 mA sa applied, PABA release was induced at pH 10. However, at pH 7, PABA was stably-bound to the chitosan-PCL matrix, with or without the external current. The selective sequestration of PABA at basic pH and its stimulated release via electric current application can be further explored for clinical translatability.

## Introduction

Controlled drug delivery systems (DDS) are smart devices regulating the sequestration, release, and bioavailability of therapeutics. Clinically-successful DDS materials include chitosan and poly(caprolactone) (PCL) as they impart safety and biocompatibility [1, 2]. Importantly, chitosan provides protonizable amine groups, enabling matrix electrostatic interaction of anionic and negatively-charged molecules [3].

Crosslinking of PCL with chitosan enables tunability of mechanical and degradation properties without affecting the DDS net charge since PCL is neutral [4–6]. Release of chitosan-bound drugs can be induced via stimulation of external electric field [7–9].

In this study, the absorption and release kinetics of *para*-aminobenzoic acid (PABA) to and from crosslinked chitosan-PCL scaffold, respectively, was characterized under different pH conditions. PABA, a benzene ring with para (carbon 1 and 4) positions of amine and carboxylic acid groups, is a zwitterion that changes in charge depending on the buffer pH. Due to its multiple pKa values [10] as well as its complex interaction with the charged chitosan-PCL matrix, PABA’s dynamic behavior is challenging to model, hence experimental outcome is needed. Moreover, PABA release was quantified in the presence of sub-milliampere electric current for the implementation of a controlled DDS with medicinal implications [11, 12].

## Materials and Methods

### Reagents

*para*-aminobenzoic acid (PABA), poly(caprolactone) (PCL; M_n_ ~ 80000), N,N’-dicyclohexylcarbodiimide (DCC), dibutyltin dilaurate (DBTDL), 4-(dimethylamino)cinnamaldehyde (DMACA), sodium acetate, 4-morpholinepropanesulfonic acid (MOPS), sodium tetraborate decahydrate (sodium borate), acetic acid, and NaOH were purchased from Sigma-Aldrich (St. Louis, MO). Chitosan (70% deacetylated) was obtained from Primex (Siglufjordur, Iceland). Deionized water (18.2 MΩ·cm resistivity) was used as the solvent, unless stated otherwise. Three buffers at different pH were prepared: 81 mM sodium acetate (pH 4), 100 mM MOPS (pH 7), and 50 mM sodium borate (pH 10).

### Chitosan-PCL hydrogel fabrication

Solutes: chitosan (40 mg/mL) and PCL (10 mg/mL) and crosslinkers: DCC (1 mg/mL) and DBTDL (0.07 μL/mL) were dissolved and reacted in 80% acetic acid at 37 °C (modified from Aroguz *et al*. [4]) in 24-well microplates (1 mL / well). Water was evaporated inside the fume hood. Scaffolds were soaked and rinsed in 1 M NaOH for two days to stabilize, then in multiple water washes until pH 7. Crosslinked chitosan-PCL were equilibrated in their respective buffers (pH 4, 7, or 10).

### PABA absorption and release

1 mg/mL PABA in pH buffers were added to their corresponding equilibrated chitosan-PCL for 2 hours to allow for PABA absorption into the bulk. The unabsorbed liquid was collected for indirect measurement of the gel-absorbed PABA (absorbed = mass added – mass unabsorbed). Gels were quickly rinsed in buffers to remove any residual unabsorbed PABA. 1 mL buffer was added at 0.5, 1, 2, and 4 hours, then the fluid was aspirated out and collected, and the scaffold was replenished with fresh 1 mL buffer. Release groups containing no absorbed PABA were used as controls.

Combined chitosan and PABA levels from the collected samples were quantified via a colorimetric reaction with DMACA (amine detection [13]) and absorbance read at 490 nm (for pH 4 and 7) and at 595 nm (for pH 10) using a spectrophotometer (iMark Reader, Bio-Rad, Hercules, CA). Chitosan release was obtained using the no-PABA (negative) controls. PABA quantities were calculated by subtracting the chitosan from the combined chitosan and PABA values.

### Electric current application

The electrical system was assembled based on Sheen and de Guzman using a MATLAB (MathWorks, Natick, MA)-controlled Arduino [14]. Briefly, electrodes spaced 1.5 cm apart (across the gel) were inserted into the scaffolds with absorbed PABA in buffers pH 7 and 10. A continuous current of 0.6 mA was applied, and liquids were collected at 0, 2, 6, 10, 14, and 18-minute intervals. Fresh buffer was added per time point. Controls were tested at no applied electric current.

### Statistical analyses

Experimental samples were performed in triplicates (n = 3). Values were reported as average ± 1 standard deviation. Saturation regression equations were fitted using least-squares method in MATLAB:

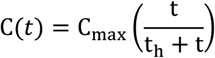

where, t = time [hr], C = cumulative mass release concentration, C_max_ = theoretical maximum concentration, and t_h_ = time to reach half C_max_. Scatterplots and one-way analysis of variance (ANOVA) with multiple comparison (Tukey’s) were made in MATLAB at 5% probability (p) of type I error.

## Results

### Chitosan-PCL gels

The fabricated chitosan-PCL scaffolds were stable in water at pH 7 and initially after equilibration with the different buffers (Fig. 1a). However, after hours of incubation, those in acidic medium (pH 4) appeared to degrade faster relative to gels in higher pH values. DMACA colorimetric assay confirmed that the amine-containing chitosan chains were released most in the acidic buffer.

**Figure 1.**
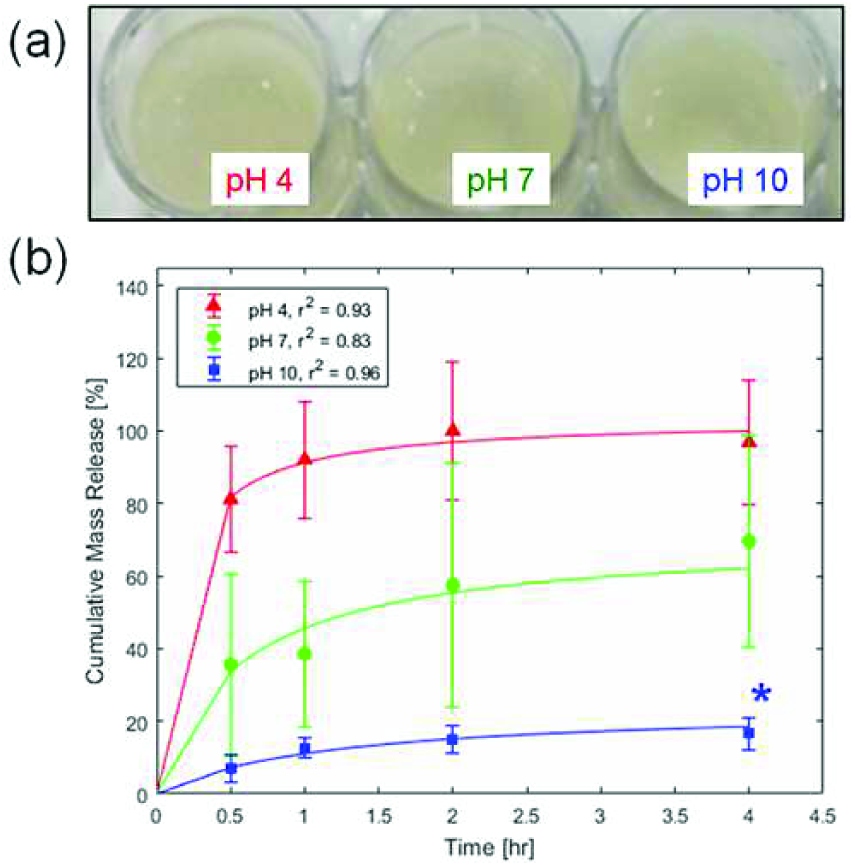
(a) Chitosan-PCL gels absorbed with PABA displayed (b) different saturation release kinetics of PABA in buffers with pH of 4, 7, and 10. *p ≤ 0.0291 compared to the other groups.

### PABA release kinetics

Chitosan-PCL gels absorbed 0.26, 0.82, and 1.73 mg PABA, at pH 4, 7, and 10, respectively. The cumulative mass release of PABA normalized to these initial masses produced saturation curves (Fig. 1b) with the fitted least-squares regression equation, 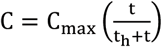 constants: C_max_ = 1.0328 and t_h_ = 0.1320 hr (r^2^ = 0.93) at pH 4, C_max_ = 0.7060 and t_h_ = 0.548 hr (r^2^ = 0.83) at pH 7, and C_max_ = 0.2387 and t_h_ = 1.1565 hr (r^2^ = 0.96) at pH 10. The t_h_ (time to achieve half of C_max_) values showed increasing trend as pH increased. Based on the saturation functions at the 4-hour time point, the amount of PABA released into the liquid media were 100% at pH 4, 62.1% at pH 7, and 18.5% at pH 10 (p ≤ 0.0291; significantly lower release compared to pH 4 and 7).

### PABA release with electric current induction

The chitosan-PCL gel system (Fig. 2a), in different pH buffers, with electrode separation distance of 1.5 cm generated electrical resistance values in the 8 kΩ range. Consequently, the highest electrical current generated with the 5-V potential of the Arduino controller was at 0.6 mA. Application of this continuous current for 18 minutes led to different PABA release profiles (Fig. 2b). At the neutral pH of 7, the release kinetics of PABA, with and without electric current were statistically similar (p ≥ 0.6479) at all time points. The longer the time, the more mass was freed from the chitosan-PCL bulk into the medium; and by 14 and 18 minutes, the release was statistically-greater (p ≤ 0.0028) relative to the control group without electric current application.

**Figure 2.**
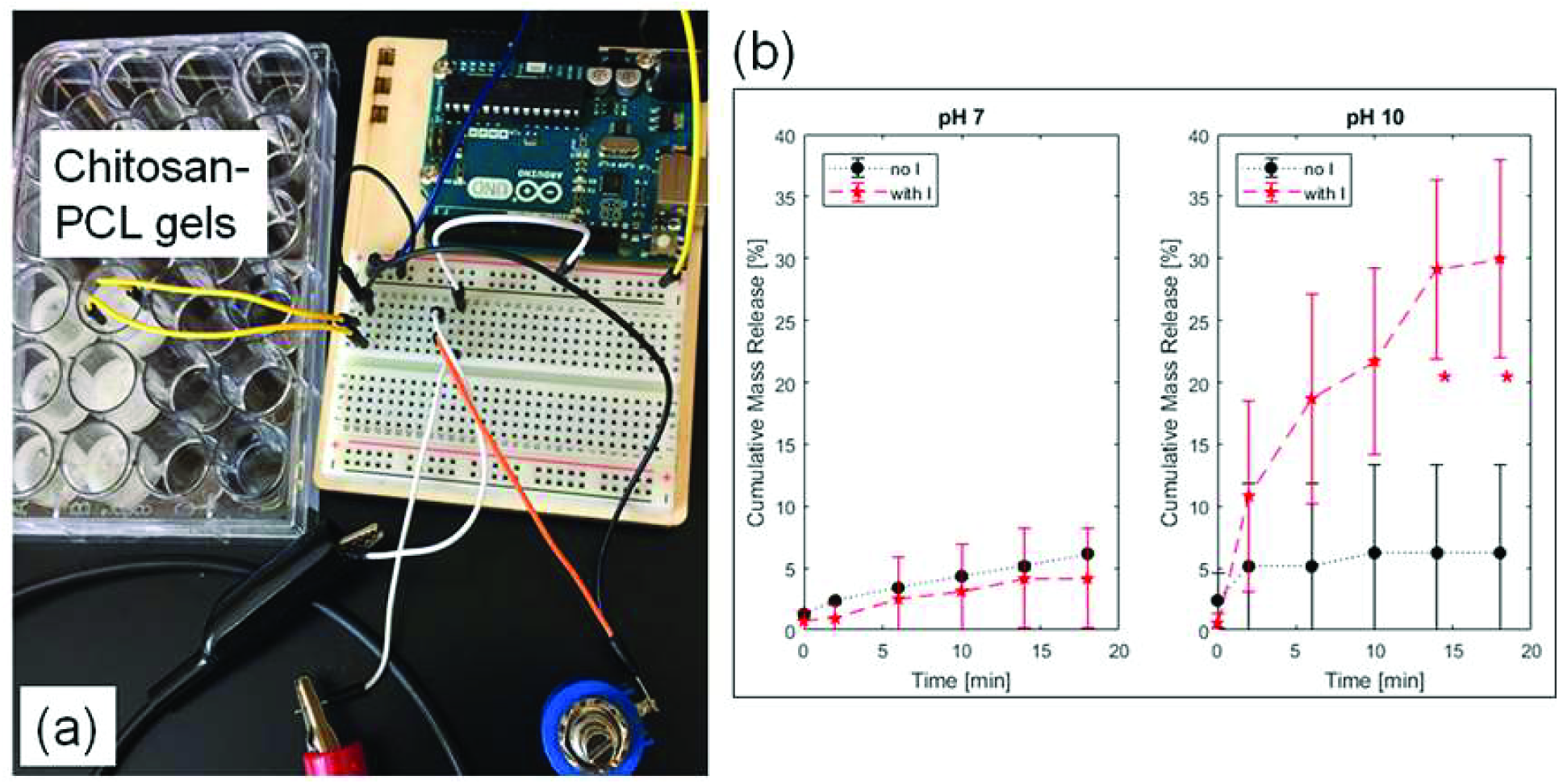
(a) Experimental setup for 0.6 mA electric current delivery (with I) into the chitosan-PCL gels to (b) release PABA. Control gels without current (no I) were compared side-by-side at pH 7 and 10. *p ≤ 0.0028 compared to the no I group.

## Discussion

The crosslinking conditions (such as solvent type and concentration, crosslinker amount, and reaction times) using DCC and DBTDL with chitosan and PCL chains can be improved to fabricate a more stable gel network that can be utilized to deliver drugs. In our current system, the unbound chitosan affected the PABA absorption and release kinetics; hence, increased datapoint variability (Fig. 1b and 2b).

Experimental results showed (Fig. 1b) that the higher the pH, the more PABA associated with the chitosan network suggesting electrostatic interaction effects. At basic pH, the positively-charged chitosan sequestered the highly negatively-charged PABA with acidic isoelectric point [15]. The PABA diffusion out of the chitosan-PCL network represented 78% to 100% total release relative to the C_max_ (theoretical maximum asymptote) values indicating that, at the neutral and basic pH media, the network-bound PABA was stable and cannot be dissociated by diffusion-mediated forces. The complete release of PABA in the acidic pH environment may be attributed to the repulsion between the highly positively-charged chitosan network and the positively-charged protonated amine group of PABA.

In the presence of an external current (Fig. 2b), the PABA release at pH 7 was found to be similar to the “no current” control. This observation suggests that PABA was neutral, thus negative charges from the electric current did not alter its mass release. However, at pH 10, the release behavior of PABA was found to be time-dependent. A possible mechanism is that, at pH 10, flowing electrons displaced the negatively-charged PABA from the positive chitosan-based matrix [8] leading to induced PABA release via electroosmosis and diffusion. This electroresponsive approach [9] can be a useful controlled drug delivery strategy.

## Conclusion

The amphoteric PABA is shown to have altered interactions with chitosan-PCL scaffolds based on pH values of the liquid media. More PABA is absorbed but less is released at higher pH. Electric current induces the release of tightly-bound PABA (at basic pH), likely due to electrostatic competition of negatively-charged PABA and flowing electrons. This controlled response can be utilized for the development of targeted PABA therapeutics and nutritional usage [11, 12].

## Acknowledgements

We would like to thank the members of the Bioengineering Materials Lab for their technical assistance, manuscript proofreading, and discussion, Jacqueline Scarola and Lori Castoria for help with purchasing, and Sina Rabbany for his continuing support and collaboration in conducting biomaterials research at Hofstra SEAS. Support for this study was provided by Hofstra University internal research funds.

## Abbreviations

DDS: drug delivery systems
PCL: poly(caprolactone)
PABA: *para*-aminobenzoic acid
DCC: N,N’-dicyclohexylcarbodiimide
DBTDL: dibutyltin dilaurate
DMACA: 4-(dimethylamino)cinnamaldehyde
MOPS: 4-morpholinepropanesulfonic acid

